# Placenta DNA Methylation Adaptation to Maternal Glucose Tolerance in Pregnancy

**DOI:** 10.1101/224139

**Authors:** Andres Cardenas, Valerie Gagné-Ouellet, Catherine Allard, Diane Brisson, Patrice Perron, Luigi Bouchard, Marie-France Hivert

## Abstract

Maternal hyperglycemia during pregnancy is associated with fetal growth and adverse perinatal and developmental outcomes. Placental epigenetic maladaptation may underlie these associations. We performed an epigenome-wide association study of term placentas and prenatal maternal glucose response 2-hour post oral glucose challenge at 24-30 weeks of gestation among 448 mother-infant pairs. Maternal glucose levels post-load were strongly associated with lower DNA methylation of 4 CpGs (FDR q<0.05) within the Phosphodiesterase 4B gene (*PDE4B*). Additionally, three other CpGs were differentially methylated relative to maternal glucose response within the *TNFRSF1B; LDLR*; and *BLM* genes (FDR q<0.05). Methylation levels correlated with expression in placental tissue for all 4 CpG sites in *PDE4B* (*r_s_*: 0.26–0.35, P<0.01), *LDLR* (*r_s_*: 0.22, P=0.03) and at *TNFRSF1B* (*r_s_*: -0.25, P=0.01). Our study provides evidence that maternal glucose response during pregnancy is associated with DNA methylation of genes within the placenta that are partially under epigenetic control.

## INTRODUCTION

Prenatal nutritional, behavioral and environmental conditions play a key role in fetal development by modulating the intra-uterine environment, fetal nutrient availability, and growth. The Pedersen hypothesis states that prenatal maternal glucose, that easily crosses the placenta, might lead to intrauterine hyperglycemia affecting fetal growth and development^1^.

It is now well recognized that maternal hyperglycemia in pregnancy is associated with adverse maternal and birth outcomes. For example, in a large prospective international multicenter study, the Hyperglycemia and Adverse Pregnancy Outcome (HAPO) study, robust linear associations were found between greater maternal prenatal glucose levels and higher birthweight or cord blood C-peptide levels across the whole spectrum of maternal glycemia, with no lower threshold^2^. In the HAPO study, measures of maternal prenatal glycemia, both fasting and post oral glucose load, were linearly associated with adverse outcomes in mothers and in infants at birth^2^. Of these, maternal glucose levels 2-hours (2h) post 75-gram glucose load showed slightly stronger associations with neonatal hypoglycemia^3^ and with an increased risk of abnormal glucose tolerance in offspring at 7 years of age^4^, suggesting that maternal response to glucose loading in pregnancy may better reflect offspring metabolic programming at birth and later in life. Indeed, emerging evidence suggests that maternal hyperglycemia, could have lasting health consequences due to metabolic programming that occurs during key fetal developmental stages and hypothesized to act through epigenetic programming and molecular mechanisms^5^.

It is now appreciated that the placenta is not simply a transferring organ for nutrients from the mother to fetus but participates actively on maternal metabolism and likely contributed to fetal programming. The placenta is the master regulator of the fetal environment and directly responsible for maternal-fetal nutrient and waste transport as well as hormone synthesis^6,7^. Fetal gluconeogenesis is relatively minimal and thus fetus relies heavily on circulating maternal glucose transported into the fetal side by facilitated placental diffusion through members of the glucose transporter (GLUT) family^6^. Over 95% of the fetal glucose levels are estimated to be derived from maternal plasma levels by diffusion^8,9^. Contrary to maternal plasma glucose, insulin does not cross the placental barrier. Instead, the fetal pancreas responds to elevated glucose by producing insulin which might lead to fetal hyperinsulinemia linked to macrosomia and increased adiposity^10,11^.

The placenta has been shown to adapt to nutrient availability through epigenomic modifications in response to gestational diabetes^12–14^. Nevertheless, these studies reported epigenetic modifications with cases of gestational diabetes, thus could not differentiate if the observed patterns were consequences of gestational diabetes (or of its treatments), and not actual placental adaptations to hyperglycemia in pregnancy. To address this gap of the role of placental maladaptation to maternal glucose levels during fetal development, we conducted an epigenome-wide association study (EWAS) for maternal 2h-glucose post oral glucose tolerance testing (OGTT) administered during the second trimester of pregnancy and DNA methylation of full term placenta tissue collected at birth. We measured genome-wide DNA methylation of the fetal placenta side among 448 mother-infant pairs and observed strong associations for several inflammatory relevant genes. Furthermore, we confirmed the functional role of DNA methylation at the discovered CpG loci by quantifying gene expression in placental tissue within our study population.

## RESULTS

### Participants’ Characteristics

A total of 448 mother-infant pairs were eligible for analyses with complete DNA methylation and 2h-glucose levels post-OGTT. All mothers were Caucasian, 49.3% were primiparous, with a mean (SD) age of 28.2 (4.3) years, mean BMI of 25.45 (5.7) Kg/m^2^ and 90.2% reported to be non-smokers during pregnancy. Mean (SD) gestational age at birth was 39.5 (1.0) weeks, mean birthweight was 3,448 (428) grams and 52.7% of births were males. At enrollment and by design of the study, none of the pregnant women had pre-gestational diabetes evaluated by first trimester glycemic testing (Glucose Challenge Testing and HbA1c). At the second trimester 75g-OGTT, mean (SD) fasting glucose level was 4.20 (0.37) mMol/L, 1h-glucose was 7.11 (1.61) mMol/L and 2h-glucose was 5.80 (1.33) mMol/L. Additional sample characteristics are summarized in **Table 1**.

**Table 1.** Sample characteristics for the Genetics of Glucose regulation in Gestation and Growth (Gen3G) birth cohort

| **Sample characteristic (N=448)** | **Mean (SD) or n (%)** |
| --- | --- |
| **Maternal age** (years) | 28.2 (4.3) |
| **BMI** (kg/m<sup>2</sup>) | 25.45 (5.7) |
| **Parity** | 2.3 (1.5) |
| Primiparous | 221 (49.3%) |
| Multiparous | 227 (50.7%) |
| **Ethnicity** |  |
| Caucasian | 448 (100%) |
| **Maternal smoking during pregnancy** |  |
| No | 404 (90.2%) |
| Yes | 39 (8.7%) |
| Unknown | 5 (1.1%) |
| **1<sup>st</sup> trimester Glucose Challenge Test (GCT)** |  |
| Glucose concentration post 50-gram challenge (mMol/L) | 5.55 (1.41) |
| **Fasting 2<sup>nd</sup> trimester Oral Glucose Tolerance Test (OGTT)** |  |
| Glucose concentration prior to 75-gram challenge (mMol/L) | 4.20 (0.38) |
| 1h-glucose concentration post 75-gram challenge (mMol/L) | 7.11 (1.61) |
| 2h-glucose concentration post 75-gram challenge (mMol/L) | 5.80 (1.33) |
| **Child gender** |  |
| Male | 236 (52.7%) |
| Female | 212 (47.3%) |
| **Gestational age at birth** (weeks) | 39.5 (1.0) |
| **Birthweight** (grams) | 3,448 (428) |

### Association of Maternal 2h-Glucose Levels with DNA Methylation of Placenta

In CpG-by-CpG analyses adjusted for maternal age, maternal BMI, parity, maternal smoking in pregnancy, child gender, gestational age at birth and bioinformatic adjustment of cellular heterogeneity, we found 7 CpG sites of placenta DNA significantly associated (FDR q<0.05) with maternal 2h-glucose levels post 75g-OGTT, **Figure 1**. We observed lower placental DNA methylation of four CpG sites annotated to *PDE4B* (Phosphodiesterase 4B gene) in response to higher maternal 2h-glucose levels. Specifically, a one mMol/L greater 2h-glucose level was associated with a 1.16%, 0.88%, 1.86% and 0.58% lower DNA methylation at cg07734160 (*P*=1.20x10^-9^), cg13866577 (*P*=1.11x10^-7^), cg03442467 (*P*=2.84x10^-7^) and cg13349623 (*P*=2.06x10^-9^), respectively. The four CpG sites were annotated within the transcription start site (1500-200 bps), first exon, or body of *PDE4B* and surrounded by another four sites reaching unadjusted statistical significance (*P*<1x10^-4^), **Figure 2**. In addition, a one mMol/L greater maternal 2h-glucose level was associated with 1.22% higher DNA methylation of cg26189983 annotated to *TNFRSF1B/TNFR2* (Tumor Necrosis Factor Receptor Superfamily Member 1B) and located within the gene body (*P*=1.70x10^-7^). We also observed an inverse association for cg20254265 annotated to Exon boundary of *BLM* (Bloom Syndrome RecQ Like Helicase gene) with a one mMol/L greater 2h-glucose level associated with a 0.63% lower DNA methylation (*P*=7.58x10^-7^). Lastly, we observed 0.27% lower DNA methylation at cg08483713 per one mMol/L greater 2h-glucose level and this CpG site was annotated to the gene body of *LDLR* (Low Density Lipoprotein Receptor gene).

**Figure 1.**
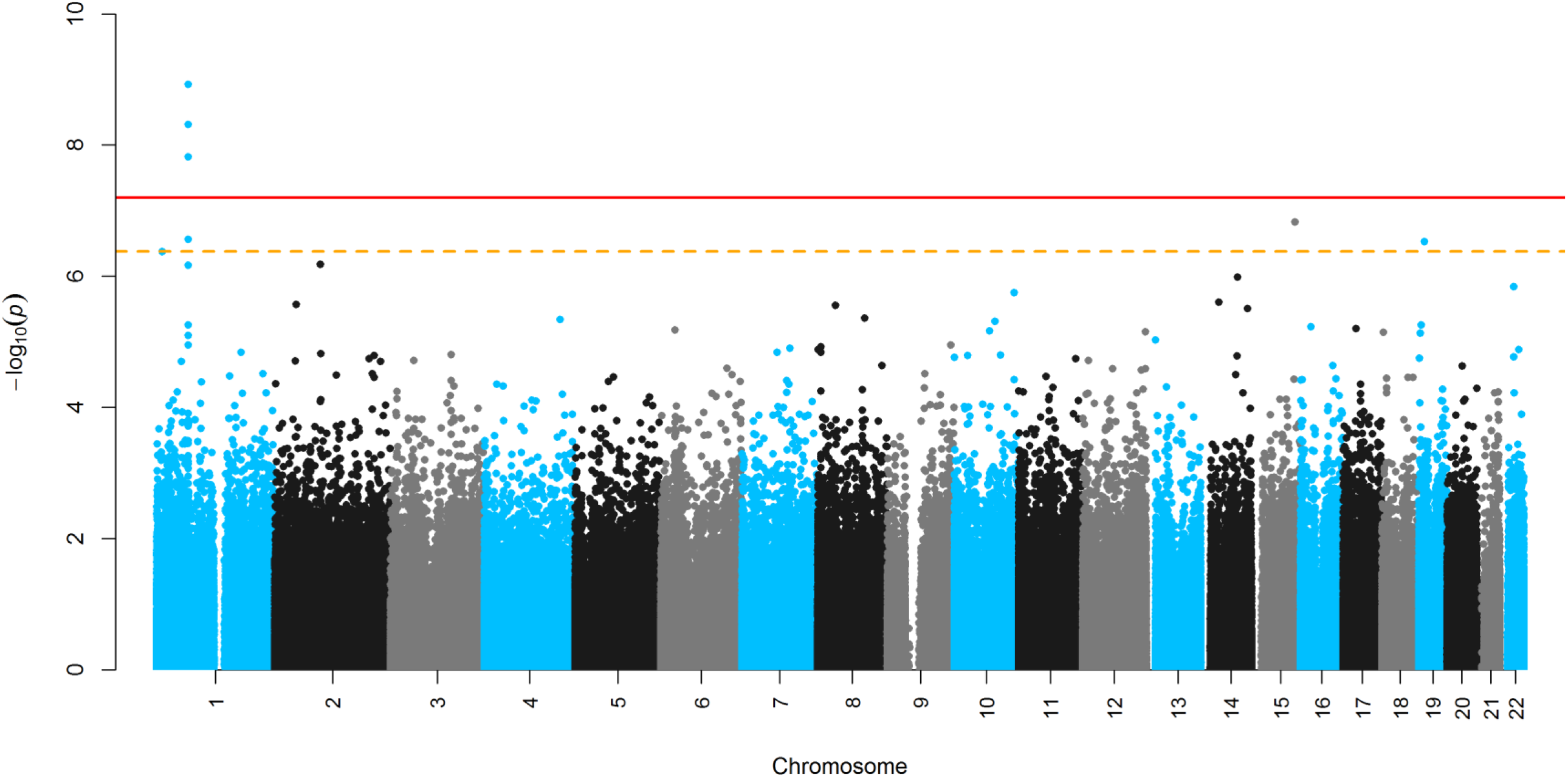
Manhattan plot for the Epigenome-wide association study (EWAS) of prenatal maternal 2h-glucose levels post OGTT

**Figure 2.**
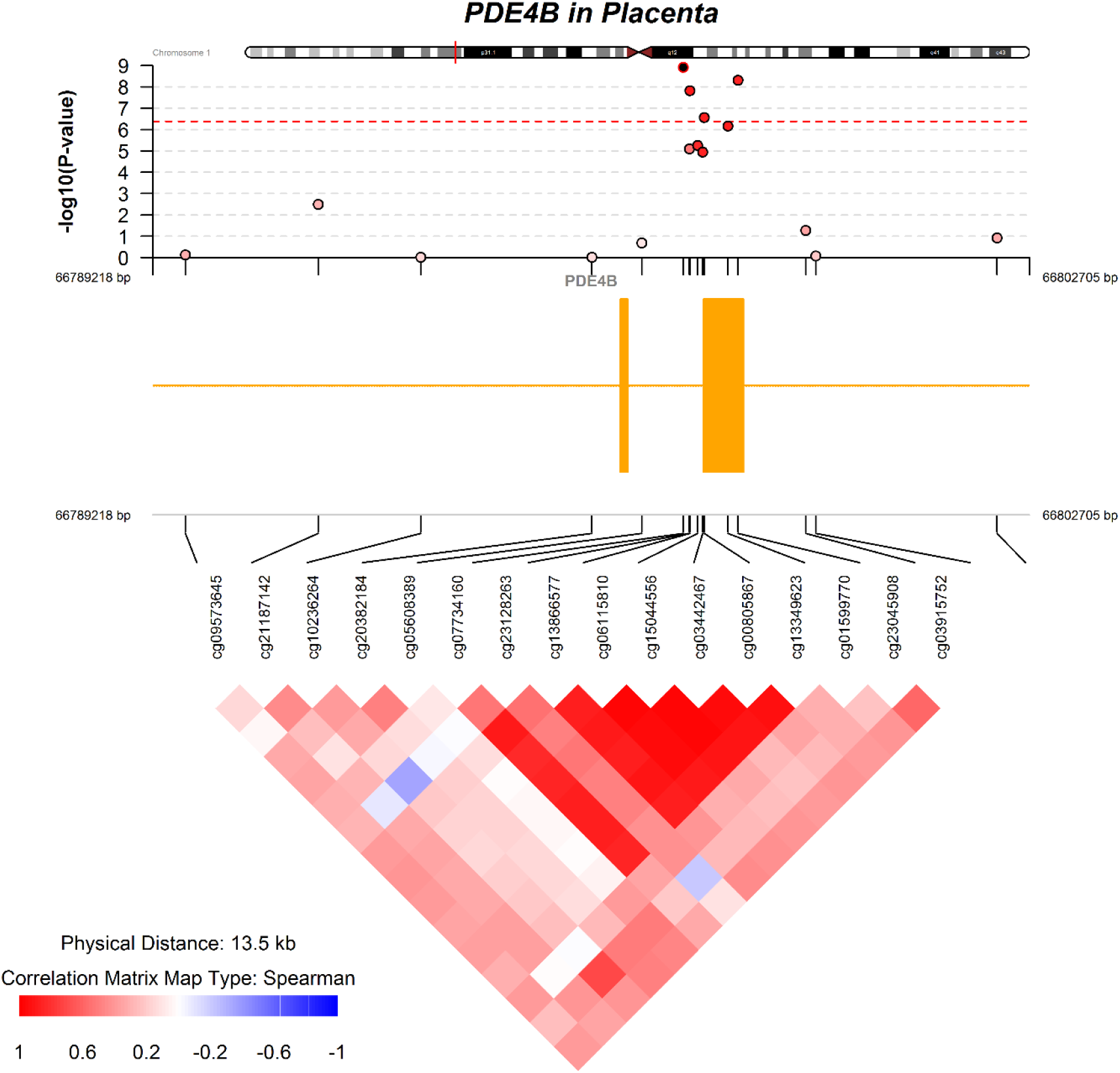
Regional Manhattan plot and correlation heatmap for CpG sites near the differentially methylated loci in the *PDE4B* gene: associations with maternal 2h-glucose levels post OGTT.

In the epigenome-wide association analyses bioinformatic adjustment reduced the genomic inflation (λ) from 1.39 to 1.01 reducing the potential impact of cellular heterogeneity on our analyses (**Figure S1**). Mean methylation levels for each CpG, genomic annotation and summary statistics for the association are presented in **Table 2** as well as their correlation distribution in **Figure S2**.

**Table 2.** Adjusted difference in DNA methylation associated with a one mMol/L increase in prenatal 2h-glucose levels post-OGTT

| CpG | Mean DNAm (SD) | Chromosome | Position | Gene | Gene group | %-difference in DNA methylation | 95% CI | P |
| --- | --- | --- | --- | --- | --- | --- | --- | --- |
| cg26189983 | 49.7% (8.2) | chr1 | 12227700 | *TNFRSF1B* | Body | 1.22% | (0.80, 1.7) | 1.70x10<sup>-7</sup> |
| cg07734160 | 13.0% (7.5) | chr1 | 66797378 | *PDE4B* | TSS1500; Body | -1.16% | (-1.5, -0.8) | 1.20x10<sup>-9</sup> |
| cg13866577 | 9.4% (6.6) | chr1 | 66797481 | *PDE4B* | Body; TSS1500 | -0.88% | (-1.2, -0.6) | 1.11x10<sup>-7</sup> |
| cg03442467 | 24.6% (13.1) | chr1 | 66797701 | *PDE4B* | Body; TSS200 | -1.86% | (-2.6, -1.2) | 2.84x10<sup>-7</sup> |
| cg13349623 | 5.8% (3.6) | chr1 | 66798221 | *PDE4B* | 1<sup>st</sup> Exon; Body | -0.58% | (-0.8, -0.4) | 2.06x10<sup>-9</sup> |
| cg20254265 | 79.6% (9.1) | chr15 | 91306178 | *BLM* | ExonBnd; Body | -0.63% | (-0.9, -0.4) | 7.58x10<sup>-7</sup> |
| cg08483713 | 5.86% (2.9) | chr19 | 11241669 | *LDLR* | Body | -0.27% | (-0.4, -0.2) | 1.39x10<sup>-6</sup> |

**Abbreviations:** TSS200= 0-200 bases upstream of the transcriptional start site (TSS); TSS1500= 200–1500 bases upstream of the
TSS; **Body**=between the ATG and stop codon; irrespective of the presence of introns, exons, TSS, or promoters; **ExonBnd**= within 20
bases of an exon boundary, i.e. the start or end of an exon.

### Association of Additional Maternal Glucose Measures with CpG sites Associated with 2h-Glucose

As sensitivity analyses, we also examined associations among differentially methylated CpG loci significantly associated with maternal 2h-glucose level post OGTT and other prenatal maternal glycemic indices in pregnancy. Namely, we tested associations with non-fasting glucose levels measured during the first trimester 1-hour (1-h) post 50g Glucose Challenge Test (GCT) as well as fasting and glucose levels 1-h post 75g OGTT measured during the second trimester. Greater post load maternal glucose at either first (1h-glucose level post 50g-GCT) or second trimester (1h-glucose level post 75g-OGTT) were associated with lower DNA methylation at previously discovered CpG sites in *PDE4B, BLM* and *LDLR* genes (*P*<0.05), consistent in the direction of association found in discovery associations with 2h-glucose level at second trimester, but with relatively smaller effect sizes.

Associations between maternal fasting glucose at second trimester and DNA methylation at CpG sites in *PDE4B* were in the same direction than initial findings in the 2h-glucose analyses but not significant. We did not find consistent associations in direction of effect first trimester 1-h post 50g or second trimester fasting glucose level and DNA methylation at *TNFRSF1B* CpG site. Associations among top differentially methylated loci discovered and other maternal glucose measures are summarized in **Table 3**.

**Table 3.**
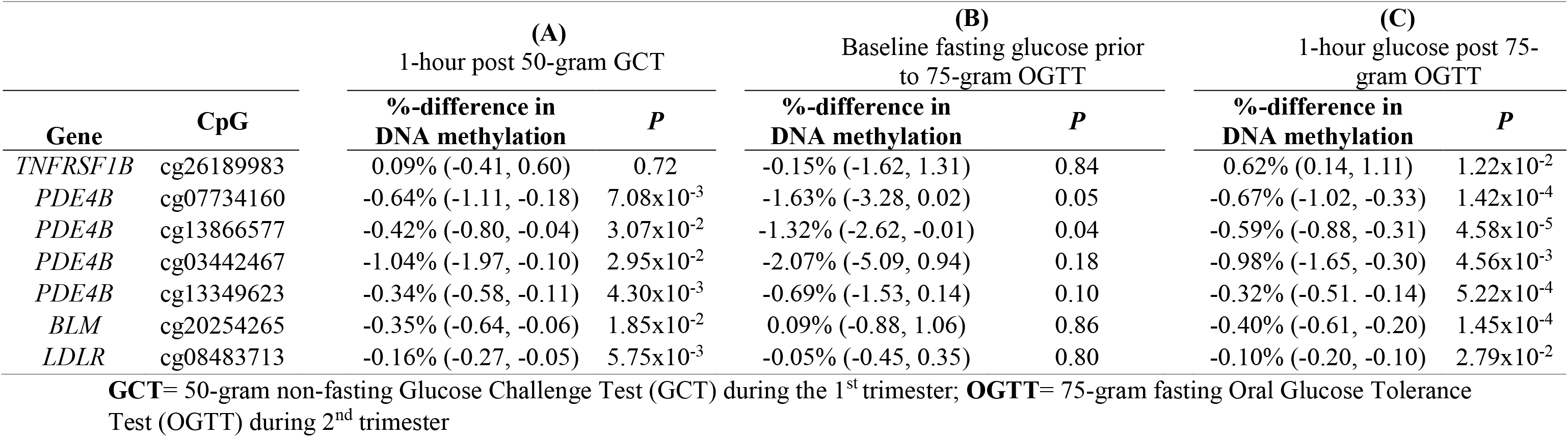
Adjusted difference in DNA methylation among CpGs discovered in the EWAS of 2h-glucose levels: A) first trimester nonfasting post 50-gram GCT glucose concentrations B) second trimester baseline fasting glucose concentrations and C) 2h-glucose levels post 75 gram OGTT glucose concentrations during the second trimester

| Gene | CpG | (A)<br>1-hour post 50-gram GCT |  | (B)<br>Baseline fasting glucose prior<br>to 75-gram OGTT |  | (C)<br>1-hour glucose post 75-gram OGTT |  |
| --- | --- | --- | --- | --- | --- | --- | --- |
|  |  | %-difference in<br>DNA methylation | *P* | %-difference in<br>DNA methylation | *P* | %-difference in<br>DNA methylation | *P* |
| *TNFRSF1B* | cg26189983 | 0.09% (-0.41, 0.60) | 0.72 | -0.15% (-1.62, 1.31) | 0.84 | 0.62% (0.14, 1.11) | 1.22x10<sup>-2</sup> |
| *PDE4B* | cg07734160 | -0.64% (-1.11, -0.18) | 7.08x10<sup>-3</sup> | -1.63% (-3.28, 0.02) | 0.05 | -0.67% (-1.02, -0.33) | 1.42x10<sup>-4</sup> |
| *PDE4B* | cg13866577 | -0.42% (-0.80, -0.04) | 3.07x10<sup>-2</sup> | -1.32% (-2.62, -0.01) | 0.04 | -0.59% (-0.88, -0.31) | 4.58x10<sup>-5</sup> |
| *PDE4B* | cg03442467 | -1.04% (-1.97, -0.10) | 2.95x10<sup>-2</sup> | -2.07% (-5.09, 0.94) | 0.18 | -0.98% (-1.65, -0.30) | 4.56x10<sup>-3</sup> |
| *PDE4B* | cg13349623 | -0.34% (-0.58, -0.11) | 4.30x10<sup>-3</sup> | -0.69% (-1.53, 0.14) | 0.10 | -0.32% (-0.51, -0.14) | 5.22x10<sup>-4</sup> |
| *BLM* | cg20254265 | -0.35% (-0.64, -0.06) | 1.85x10<sup>-2</sup> | 0.09% (-0.88, 1.06) | 0.86 | -0.40% (-0.61, -0.20) | 1.45x10<sup>-4</sup> |
| *LDLR* | cg08483713 | -0.16% (-0.27, -0.05) | 5.75x10<sup>-3</sup> | -0.05% (-0.45, 0.35) | 0.80 | -0.10% (-0.20, -0.10) | 2.79x10<sup>-2</sup> |

**GCT**= 50-gram non-fasting Glucose Challenge Test (GCT) during the 1<sup>st</sup> trimester; **OGTT**= 75-gram fasting Oral Glucose Tolerance Test (OGTT) during 2<sup>nd</sup> trimester

### DNA Methylation and Gene Expression

Among 104 randomly selected participants, we observed that greater DNA methylation levels at all four CpG sites found within *PDE4B* were significantly correlated with higher *PDE4B* expression in placental tissue with the strongest association observed for cg03442467 (*r_s_*= 0.35; *P*=2.4x10^-3^), **Figure 3**. Consistent positive expression DNA methylation associations were observed for cg07734160 (*r_s_*= 0.28; *P*=4.5x10^-3^), cg13866577 (*r_s_*= 0.26; *P*=7.2x10^-3^), and cg13349623 (*r_s_*=0.27; *P*=6.2x10^-3^) of *PDE4B*, **Figure S3**. Greater DNA methylation at the *LDLR* CpG site was associated with greater *LDLR* expression (*r_s_*=0.22, *P*=0.03), whereas greater DNA methylation at the *TNFRSF1B* CpG site was associated with lower TNFRSF1B expression (*r_s_*= - 0.25, *P*=0.01) of placenta tissue. We found no evidence of associations between *BLM* DNA methylation and expression at the discovered site (*r_s_*= -0.09, *P*=0.37), **Figure 3**.

**Figure 3.**
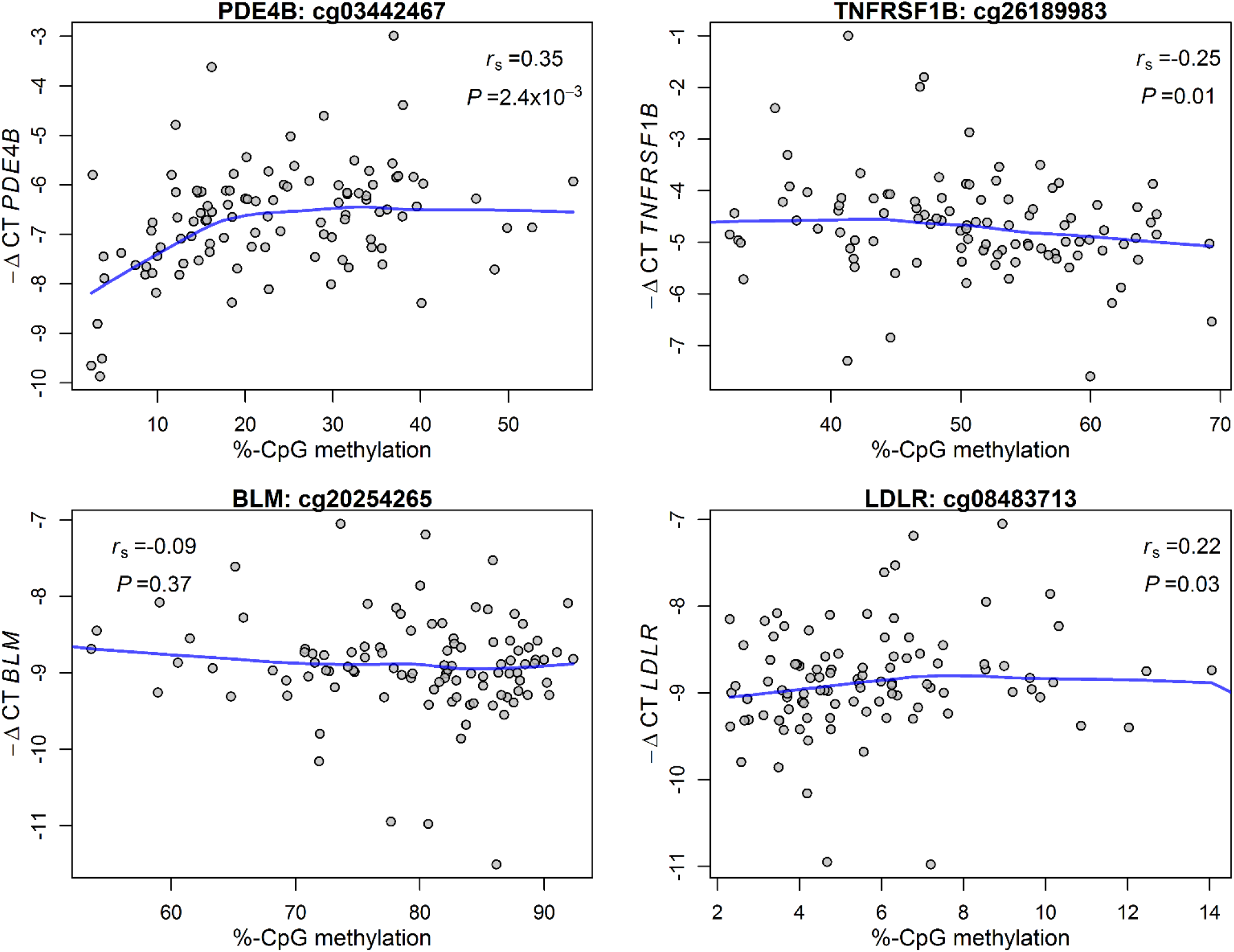
Spearman correlations coefficients and locally weighted scatterplot smoothing lines for the association between placenta DNA methylation and gene expression (-ΔCT) among loci associated with prenatal maternal 2h-glucose levels post OGTT (N=104).

### Replication of Results in an Independent Study Cohort: ECO-21

In an external independent birth cohort with a smaller sample size (*N*=66 to 111) we sought to replicate of our top differentially methylated site (cg26189983) at the *TNFRSF1B* gene and three other CpGs (cg07734160; cg13866577; cg03442467) within the *PDE4B* region. We did not see any significant external replication within this independent cohort (*P*>0.05).

However, estimated adjusted associations for the top discovered loci within the *TNFRSF1B* and the top CpG in the *PDE4B* region were similar in direction of association with maternal 2h-glucose levels in normoglycemic women (supplementary **Table S1**). In contrast, among ECO21 women with gestational diabetes mellitus (GDM) (n=29) the top *PDE4B* discovered CpG site, cg07734160, was positively associated with glucose 2h-glucose levels (*P*=0.01). Namely, for GDM pregnancies in ECO21 a mMol/L increase in 2h-glucose levels was associated with 4.67% higher DNA methylation at cg07734160, supplementary **Table S2**.

## DISCUSSION

In this pre-pregnancy birth cohort, we conducted the largest EWAS of placenta for maternal prenatal glucose response during pregnancy thus far. We found CpG sites for which DNA methylation levels were associated with maternal glucose response 2-hours post 75g glucose loading, performed during the second trimester of pregnancy. Notably, there was evidence that multiple CpG sites in a genomic region of the Phosphodiesterase 4B gene (*PDE4B*) were strongly and inversely associated with higher maternal 2h-glucose levels. Furthermore, greater methylation levels at identified CpG sites in the *PDE4B* locus correlated with higher gene expression of PDE4B in placental tissue, supporting the functional role of these placental epigenetic markers influenced by maternal glucose response in pregnancy. Other discovered epigenomic loci mapped to three additional genes; *TNFRSF1B, BLM* and *LDLR* observed to be associated with maternal 2h-glucose, of which DNA methylation at *TNFRSF1B* and *LDLR* loci correlated with expression of respective gene in placental tissue. Our study provides evidence that maternal glucose response post challenge in mid-pregnancy is associated with differential methylation of genes within the placenta at birth and these loci are partially under epigenetic control for gene expression. Our results highlight the ability of the placenta to epigenetically adapt to the maternal nutritional environment that might play a functional role in metabolic programming of the offspring.

The Phosphodiesterase 4B gene (*PDE4B*) is a member of the cyclic nucleotide phosphodiesterases family, responsible for the hydrolysis of cyclic AMP and GMP^15^. The PDE4 family of enzymes catalyzes the hydrolysis of second messenger cyclic AMP (cAMP), a key signaling molecule for immune response regulation^16^. Inhibition of PDE4 decreases TNFα secretion^17^, a potent pro-inflammatory cytokine^18^. Specifically, the *PDE4B* isoform has been shown to predominantly mediate TNFα release^17^. Indeed, PDE4 inhibition in mice has been shown to block intrauterine inflammation, decrease cytokine production and delay preterm birth^19^. For instance, in an experimental mouse study, intrauterine injection with *Escherichia coli* LPS increased expression of PDE4B eliciting an inflammatory response and triggering preterm delivery. In this mouse study, PDE4B inhibition blocked intrauterine inflammatory response and prevented pre-term delivery. Remarkably, this inflammatory response was localized in glycogen trophoblast cells of the placenta from the fetal compartment, suggesting a direct role of the placenta^20^. The relationship between PDE4 inhibition and the pro-inflammatory cytokine TNFα is of high interest as TNFα levels have been observed to be increased in adipose and placenta tissue of obese pregnant women compared normal weight^21^. Circulating levels of TNFα are also a strong independent predictor of insulin resistance in pregnancy with placenta tissue being a primary contributor to maternal TNFα levels^22^. The association between insulin resistance and TNFα levels in pregnancy has also been confirmed in our cohort^23^.

There is emerging evidence that suggests that PDE4B plays an important role in adiposity and metabolic function. For example, PDE4B-null mice have been shown to be leaner, with lower fat pad, smaller adipocytes, and decreased serum leptin levels compared to wild-type littermates^24^. Treatment with a PDE4 inhibitor reduced the body weight of mice fed a Western-type diet mediated by an increase in energy expenditure and PDE4B mRNA in white adipose tissue^25^. Furthermore, chronic treatment with PDE4 inhibitors was shown to delay the progression of diabetes among an experimental animal model for obesity, diabetes and metabolic syndrome (db/db mice)^26^. In addition, a randomized controlled trial of newly diagnosed patients with type 2 diabetes demonstrated that treatment with a PDE4 inhibitor (roflumilast) successfully lowered glucose levels^27^.Therefore, PDE inhibitors have been proposed as a therapeutic agent for diabetes and metabolic syndrome^28,29^. Our results add to the growing body of evidence suggesting that *PDE4B* plays a functional role in metabolic programming and that placental DNA methylation of *PDE4B* might play a key role in this relationship.

Higher DNA methylation of TNF Receptor Superfamily Member 1B (*TNFRSF1B*) loci was associated with greater 2h-glucose levels and a weaker positive association was also observed with 1h-glucose levels post load during the second trimester of pregnancy. *TNFRSF1B* encodes a high-affinity receptor for TNFα. TNFα is linked to metabolism, insulin sensitivity in human tissues as well as in experimental studies and genetically linked to hyperlipidemia^30,31^. Expression of TNFRSF1B, also known as *TNFR2*, has been detected in placental trophoblasts and distributed across the cytosol with multiple functions such as apoptosis, inhibition of trophoblast cell fusion and invasion or epithelial shedding^32^. In an animal model, pharmacological attenuation of TNF-α signaling with soluble TNFR2-IgG (Etanercept) protected the placenta of deformities due to infection. In obese adult individuals prolonged treatment with Etanercept improved fasting glucose and adiponectin levels^33^. Furthermore, among individuals with normal glucose levels plasma soluble TNFR2 (sTNFR2) was negatively associated with insulin sensitivity^34^ while higher sTNFR2 has been observed in offspring of type 2 diabetic subjects^35^. In line with these findings plasma sTNFR2 concentration has been proposed to serve as a marker of TNFα-related insulin resistance^36^. Our findings support the hypothesis that higher maternal glucose response post challenge is associated with lower DNA methylation of *TNFRSF1B* which in turn is associated with lower expression levels. Epigenomic modifications at the *TNFRSF1B* and *PDE4B* suggests a role for TNFα regulation, a pro-inflammatory cytokine amply associated with insulin sensitivity and metabolic dysrregulation^37–39^.

Another CpG site associated with prenatal maternal glucose response was annotated to the body of the Low-Density Lipoprotein Receptor (*LDLR*). The *LDLR* encodes for a lipoprotein receptor that mediates endocytosis of LDL particles into the cell and expressed in the placenta^40^. Human studies have shown that intrauterine growth restriction is associate with changes in placental LDLR expression compared to normal pregnancies^41,42^. Additionally, an increase in placenta LDLR expression has been documented in pregnancies with gestational diabetes in full-term placentas and suggested to be attributed to maternal inflammation^40^. Supporting this hypothesis, *in vitro* studies have shown that inflammatory cytokines such as TNF-α regulate cholesterol-mediated LDL receptor regulation^43^. This adds to the current body of evidence that points at differential methylation of genes associated with lipid transfer such as *LPL* associated with maternal glycemia during pregnancy^44^.

Lower DNA methylation at a CpG site annotated to the body of the Bloom Syndrome RecQ Like Helicase gene (*BLM*) was associated with greater maternal 2h-glucose level post oral glucose load both in the first and second trimesters. The *BLM* gene codes for an enzyme that restores replication breaks in DNA and is associated with genome stability and maintenance. Mutations of this gene are associated with an autosomal recessive syndrome, Bloom syndrome^45^. However, its role in glucose homeostasis or placental functions is unknown. Furthermore, this was the only loci not associated with gene expression in the placenta. Given the limited literature on *BLM* and glucose response this finding must be interpreted with caution.

Although results were not directly replicated in a smaller independent cohort of nondiabetic pregnant women, the top CpG from the *PDE4B* gene was associated with maternal 2-hour glucose in mothers with Gestational Diabetes Mellitus (GDM), yet in the opposite direction of what we had found in the discovery cohort. This finding may be due to intensive treatment of women with GDM, leading to lower maternal glucose over the third trimester. Post-load glucose is one of the main targets of gestational diabetes treatment, and we know that women with GDM in the replication cohort were intensively followed and tightly controlled. Nevertheless, our findings suggest that this site is epigenetically controlled by maternal glycemic response.

Our findings might be directly linked to prenatal maternal impaired glucose response where impaired response give rise to DNA methylation alterations reflected in placental DNA collected at birth. Alternatively, the observed associations might be part of the pathophysiology of impaired glucose response during pregnancy and therefore DNA methylation shifts are reflected as biomarker of this physiological process. Our prospective design minimizes the latter but does not rule out the possibility of reverse causality as it is not possible to know at what point these marks where established in pregnancy.

### Conclusion

In this prospective study of healthy expecting mothers and term births, we observed robust associations between maternal glucose response and DNA methylation of the placenta at several genes implicated in inflammatory processes and experimentally linked to pre-term birth. Methylation levels at the discovered loci correlated with functional changes in gene expression potentially reflecting placental adaptions to maternal impaired glucose response that could underlie fetal metabolic programming.

## METHODS

### Study Population

Samples and study participants were selected from the Genetics of Glucose regulation in Gestation and Growth (Gen3G), a large prospective Canadian pre-birth cohort. Gen3G was designed to elucidate the biological, environmental, genetic and epigenetic determinants of glucose regulation during pregnancy and the impact in offspring development as previously described in detail elsewhere^46^. Briefly, we recruited expecting mothers during the first trimester of their pregnancy and we enrolled pregnant women in the study if they were at least 18 years of age or older with a singleton pregnancy and did not have pre-pregnancy diabetes based on medical history and screening during the first trimester blood sampling. For this study, mother-infant pairs were selected from the larger cohort if they had placental tissue available for DNA isolation as well as >37 weeks of gestation at delivery. Study participants provided written informed consent prior to enrollment in accordance with the Declaration of Helsinki. All study protocols were approved by the ethical review board from the Center Hospitalier Universitaire de Sherbrooke (CHUS).

### Placental tissue, DNA and RNA Extraction

Trained research personnel collected fetal placenta tissue samples immediately after delivery (<30 minutes postpartum). A one cm^3^ of placenta tissue sample was collected approximately 5 cm from the umbilical cord insertion, from the fetal side of the placenta for each delivery. Placenta samples were collected by trained study staff and stored in RNALater (Qiagen, USA) at -80 ^0^C until DNA or RNA extraction occurred. We purified DNA and RNA from the placenta samples using the All Prep DNA/RNA/Protein Mini Kit (Qiagen, USA). Purity of extracted DNA was evaluated using a Spectrophotometer (Ultrospec 2000 UV/Visible; Pharmacia Biotech, USA) with an absorbance ratio set at 260 to 280 nm as recommended^47^.

### Oral Glucose Tolerance Test

During the first trimester, enrolled participants completed a non-fasting 50-gram Glucose Challenge Test (GCT) and we measure glucose 1-hour after the glucose load. During the second trimester clinical visit, all women performed a fasting 75-gram oral Glucose Tolerance Test (OGTT) and we measured maternal glucose levels were at fasting prior to the oral glucose challenge, 1-hour and 2-hour post glucose challenge. Maternal blood glucose concentrations pre- and post-OGTT were measured at a centralized hospital location.

### Epigenome-Wide DNA Methylation Measurements

Epigenome-wide DNA methylation measurements were performed on DNA extracted from the placenta samples using bisulfite conversion followed by quantification using the Infinium MethylationEPIC BeadChip (Illumina, San Diego, CA). The Infinium MethylationEPIC BeadChip measures over 850,000 CpGs at a nucleotide resolution for each sample. Samples were randomly allocated to different plates and chips to minimize confounding by technical batch effects. Methylation data was imported into *R* for preprocessing using *minfi*^48^. We first performed quality control at the sample level, excluding samples that failed (n=8), mismatch on genotype (n=12) or sex (n=1) and technical duplicates (n=10). A total of 448 high quality samples were retained for subsequent analyses. We performed quality control on individual probes by computing a detection *P*-value relative to negative control probes in the array. Namely, we excluded 2,003 probes with non-significant detection (*P*>0.05) for 5% or more of the samples. We also excluded 19,129 probes annotated to sex-chromosomes, 2,836 non-CpG probes, 5,552 SNP associated probes at the single base extension with a minor allele frequency of ≥5% and 4,453 probes with a SNP at the target CpG with a minor allele frequency of ≥5%. Finally, we excluded 40,448 cross-reactive probes previously identified^49^. A total of 791,131 CpGs were included in the final analyses. We preprocessed our data using functional normalization with the default of two principal components from the control probes^50^. We also adjusted for probe-type bias using RCP, a regression method approach that uses genomic proximity to adjust the distribution of type-2 probes^51^. Lastly, we used the ComBat function from the *sva* package to adjust for sample plate as a technical batch variable^52^. We visualized the data using density distributions at all processing steps and performed principal component analyses to examined the association of both technical and biological variables with DNA methylation variation using ENmix^53^, to ensure technical batch adjustment. We logit transform the β-values to M-values for statistical analyses as they have been shown to best meet statistical modeling assumptions^54^. However, we report effect estimated and summary statistics on the β-value scale to ease interpretability.

### RNA Quantification of Top Differentially Methylated Loci

We selected a simple random sample of 104 participants with DNA methylation data for RNA quantification. RNA concentrations and RNA Integrity Number (RIN) were assessed using Agilent 2100 Bioanalyzer and the Agilent RNA 6000 Nano Kit (Agilent Technologies, USA). Complementary DNA (cDNA) of placental RNAs were generated using a random primer hexamer (High Capacity cDNA RT, Applied Biosystems, USA). Amplicons were generated in duplicate in 20μl with 10μ of TaqMan^®^ Universial PCR Master Mix (Applied Biosystems, USA). RNA expression for genes annotate to the differentially methylated loci significantly associated with 2-hour maternal glucose (*PDE4B*: Hs00277080_m1, *TNFRSF1B*: Hs00961748_m1, *BLM*: Hs00172060_m1, *LDLR*: Hs00181192_m1, Applied Biosystems, USA) were measured with quantitative real-time PCR (qRT-PCR) using 7500 Real Time PCR system (Thermofisher, USA). Expression levels are reported relative to a reference gene, the tyrosine 3-monooxygenase/tryptophan 5-monooxygenase activation protein zeta (*YWHAZ*: Hs01122445_g1, Applied Biosystems, USA) previously shown to be stable in human placenta^55^. We report relative expression using the inverse change in the threshold cycle -ΔCT= CT (target gene) – CT (YWHAZ).

### Replication Cohort and Pyrosequencing: ECO-21

The independent replication cohort (ECO21) consisted of mother-infant pairs recruited in the first trimester of pregnancy, from a French-Canadian population from the Saguenay-Lac-Saint-Jean region of Québec (Saguenay city area, Québec, Canada)^56^. Pregnant women were excluded if they were <18 or >40 years of age, had a history of alcohol/drug abuse in pregnancy, diagnosed familial hypercholesterolemia, pre-gestational diabetes or other pre-pregnancy disorders impairing glucose homeostasis. The Chicoutimi Hospital Ethics Committee reviewed and approved this project. All women provided written informed consent prior to enrollment in the study in accordance with the Declaration of Helsinki.

In the replication cohort DNA methylation levels of the top CpG site annotated to the *TNFRSF1B* gene (cg26189983) as well as three other CpG sites (cg07734160, cg13866577 and cg03442467) within the *PDE4B* region were measured using pyrosequencing (Pyromark Q24, Qiagen, USA). Briefly, DNA underwent a sodium bisulfite (NaBis) treatment (EpiTect Bisulfite Kits; Qiagen, USA). Target NaBis-DNA loci were PCR-amplified with specific primers designed using the PyroMark Assay Design software (version 2.0.1.15; Qiagen, USA) and were then pyrosequenced. Each of the pyrosequencing runs performed included, a negative PCR control, and sodium bisulfite conversion controls. Additionally, pyrosequencing quality control (peak height, deviation from reference sequence pattern and unexpected peak height) was assessed for each sample, as recommended by the manufacturer, using Pyromark Q24 Analysis Software (v1.0.10.134).

### Statistical Analyses

Among the 448 participants eligible for analyses we report our sample’s demographic and biological characteristics using means, standard deviations, or proportions. We performed CpG-by-CpG analyses by fitting robust linear regression models for each site adjusted for covariates with DNA methylation as the response variable on the M-value scale using maternal glucose levels 2-hours post OGTT as the main predictor. To control for genomic inflation and cell-type heterogeneity we used ReFACTor, a reference-free method that adjusts for cell-type mixtures in genome-wide DNA methylation studies from heterogenous tissues^57^. Robust linear regression models were adjusted for maternal age in years, BMI, parity, smoking during pregnancy, gestational age at birth, sex and the first 10 principal components estimated from ReFACTor as proxy for placenta cellular heterogeneity.

CpG-by-CpG analyses were controlled for the false discovery rate (FDR) at 5% using q-value<0.05. Quantile-quantile plots for the regression P-values were used to visually inspect genomic inflation and we report the genomic inflation factor (λ) for unadjusted and cell-type adjusted analyses. Regional and genome-wide Manhattan plots were used to report results from the epigenome-wide association analysis.

We calculated Spearman correlation coefficients (rs) to estimate the associated between DNA methylation and gene expression among a randomly selected sample of 104-participants for top differentially methylated placenta genes associated with maternal glucose response (FDR<0.05). Significant statistical correlations between DNA methylation and gene expression were considered for Spearman correlations coefficients with *P*<0.05.

In sensitivity analyses we evaluated associations among top differentially methylated CpGs (FDR<0.05) and maternal non-fasting glucose levels 1-hour post 50-gram challenge during the first trimester of pregnancy, baseline fasting glucose levels prior to 75-gram OGTT and glucose levels 1-hour post 75-gram OGTT.

## Acknowledgments

This work was supported by Fonds de recherche du Québec en santé #20697 (M.F.H); Canadian Institute of Health Research #MOP 115071 (M.F.H); American Diabetes Association accelerator award #1-15-ACE-26 (M.F.H)

